# A genetic safeguard for eliminating target genes in synthetic probiotics in response to a loss of the permissive signal in a gut environment

**DOI:** 10.1101/2024.12.16.628630

**Authors:** Nhu Nguyen, Miaomiao Wang, Lin Li, Clement T. Y. Chan

## Abstract

Following the development of therapeutic probiotics, there is an emerging demand for constraining engineered microbial activities to ensure biosafety. Many biocontainment studies developed genetic devices that involve cell death and growth inhibition on the engineered microbes, which often create basal levels of cytotoxicity that hamper cell fitness and performance on therapeutic functions; furthermore, these toxic pathways may promote genetic instability that leads to mutations and breakdown of biocontainment circuit. To address this issue, here we explore a circuit design that destroys the engineered genetic materials in a probiotic strain, instead of killing these cells, under non-permissive conditions. Our safeguard circuit involves a two-layered transcriptional regulatory circuit to control the expression of a CRISPR system that targets the engineered genes for degradation. In *Escherichia coli Nissle 1917* (*EcN*), the biocontainment system continuously scavenged and destroyed the target until no engineered cellular function could be detected, suggesting this strategy has the potential to avoid escapee formation. Additionally, this safeguard circuit did not affect *EcN* cell fitness. We further demonstrated that the engineered probiotics inhabited in mouse guts and continued the engineered activities for at least 7 days when the permissive signal was supplied constantly; when the permissive signal was not provided, the engineered activities became undetectable within two days. Together, these studies support that our safeguard design is feasible for synthetic probiotic applications.

**HIGHLIGHTS:**

- Our safeguard system only destroys target genes and does not kill the host microbes
- It terminated engineered activities in guts in response to a loss of a signal
- This safeguard allowed synthetic probiotics to inhabit in guts for at least a week
- Cellobiose has great potential to serve as a continuous genetic signal in guts

## INTRODUCTION

Microbial engineering is an emerging approach for medical applications, which has recently been explored to develop a cure for a genetic disease (*1*). Probiotics were also engineered to detect pathogens and suppress their growth (*2-5*). Researchers have developed probiotics to secrete hormone-like molecules that may control inflammatory bowel disease, diabetes, and obesity (*6-12*). For many proposed clinical applications, engineered probiotics are expected to inhabit the gastrointestinal tract of patients and, in many cases, there is a practical need to eliminate the synthetic probiotic activities from the patient after therapy, preventing any potential long-term adverse effects. Additionally, the gut is an open system that allows these microbes to be released into the environment and get in contact with other human beings. The rapid development of engineered probiotics approaches raises concerns about the importance of biosafety measures to control engineered microbes’ persistence in patients and their spread in the natural environment and our society (*13-15*).

A broad range of biocontainment designs have been developed recently to deploy in diverse microbial hosts. These designs typically require a specific molecule to create a permissive environment, and the absence of this molecule causes cell death. Many of these kill switches involve a genetic circuit approach to detect the permissive conditions and based on that to control the expression of toxic genes for killing host microbes (*16-19*). Biocontainment systems that produce a lethal effect unavoidably reduce the fitness of their hosts, which potentially affects the growth and performance of the engineered microbes. Furthermore, background cytotoxicity from a kill switch generates an evolutionary pressure for the host to eliminate toxic genetic elements, hampering the stability of the biocontainment system (*16, 20, 21*). To enhance the stability of genetic safeguards, researchers have spent a huge amount of effort to develop various strategies, such as using multiple lethal pathways (*19, 22, 23*), blocking genetic recombination pathways (*16*), and expressing antitoxins to reduce background toxicity under permissive conditions (*17, 24*). Another strategy to provide high stability in biocontainment is synthetic auxotrophy that depends on non-standard amino acids for essential protein expression (*25-27*); however, it requires extensive genome editing, which can be challenging, especially for probiotic strains with limited genetic characterization.

Instead of killing the probiotics, in this study, we explore a strategy that uses a genetic circuit approach to only eliminate exogenous genetic materials in a non-essential plasmid, returning engineered cells to their native form, which becomes a microbial strain that naturally exists in gut microbiomes. Our engineered probiotics maintained the target plasmid in the permissive environment with a specific input signal and the loss of this input triggered the degradation of the target exogenous gene. Since the exogenous gene is orthogonal to endogenous cellular pathways, its destruction is not expected to affect fitness nor create selective pressure for inactivating our biocontainment system. Once the target plasmid is eliminated, the engineered probiotic activities are removed from the gut microbiomes.

Our design involves a CRISPR system to eliminate target genes. Previous studies demonstrate that expressing CRISPR systems can efficiently remove DNA with the targeted sequence in cellular environments (*28, 29*). With a two-layered inducible transcriptional repression circuit to control the CRISPR system (**Figure 1A**), the permissive signal is required to repress *CRISPR* gene expression.

**Figure 1.**
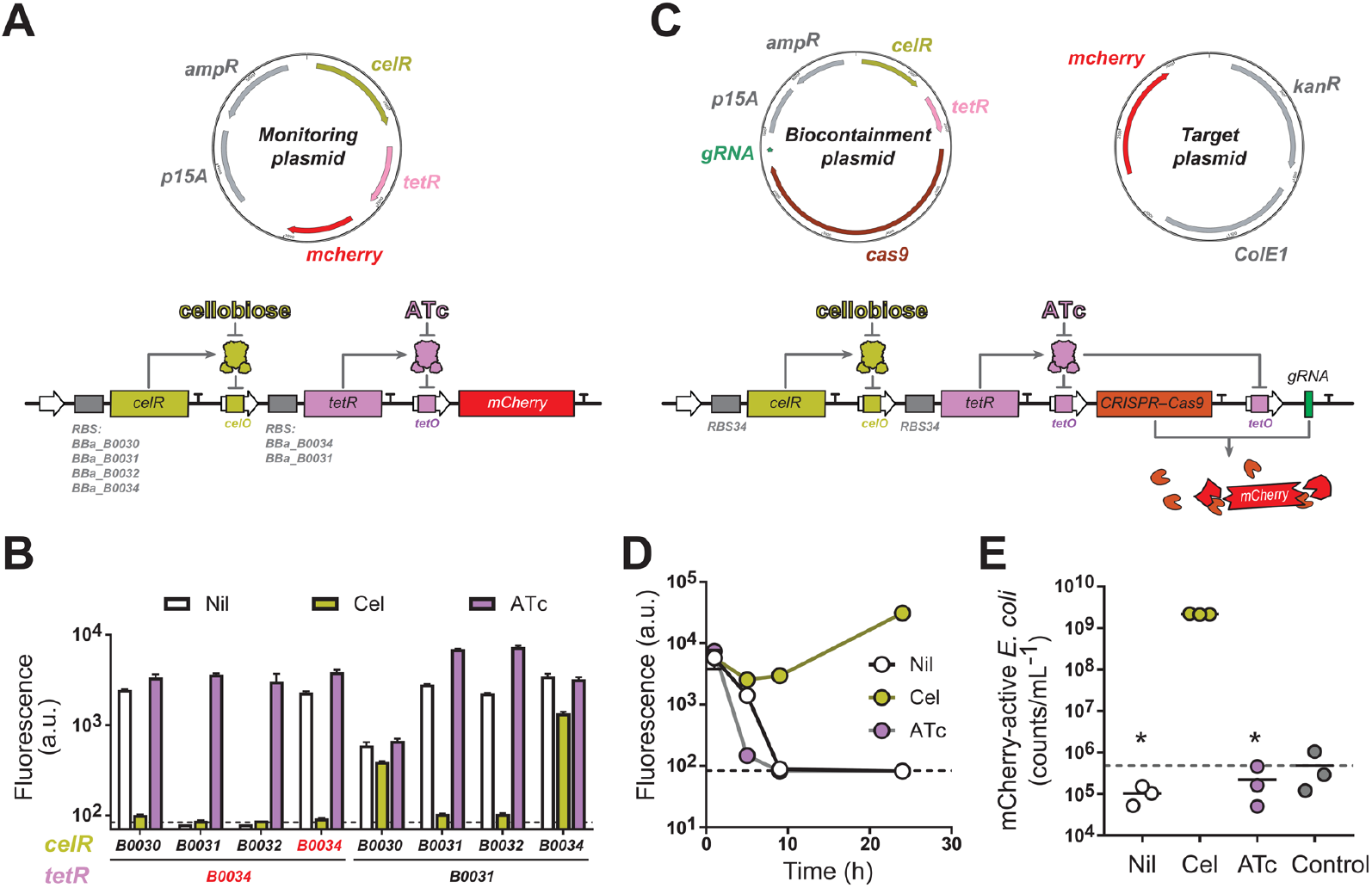
Development of the biocontainment system in *E. coli K-12*. (**A**) The design of circuit for permissive signal-monitoring. The circuit was incorporated into a plasmid with an ampicillin-resistance cassette and a *p15A* origin of replication. Eight versions of this circuit were constructed with different DNA fragments to serve as ribosomal binding sites for *celR* and *tetR*. (**B**) Characterization of permissive signal-monitoring circuits. *E. coli* cells containing each version of the circuit were exposed to no inducer (Nil), cellobiose (Cel) or anhydrotetracycline (ATc), and their levels of mCherry fluorescence were measured with flow cytometry. RBSs selected for the final version of the monitoring system are colored in red. In panels **B** and **D**, each data point represents the mean ± S.D. of three biological replicates; the error bar is not shown if it is shorter than the marker. In panels **B, D**, and **E**, the average value from three control samples, from wild-type *E. coli* cells, is presented with the dotted line. (**C**) The design of biocontainment circuit for degrading target genetic materials. Our engineered cells hosted two plasmids; one contained the biocontainment system with an ampicillin-resistance cassette and a *p15A* origin of replication (see **Supplementary Data 1**) and the other plasmid, as the target for destruction, contained the constitutively expressed *mcherry* gene, a kanamycin-resistance cassette, and a *ColE1* origin of replication (see **Supplementary Data 2**). (**D**) Characterization of target exogenous gene expression activities of *E. coli* with the biocontainment system in permissive and non-permissive conditions. The geometric mean of mCherry fluorescence in the engineered *E. coli* population was measured with flow cytometry at 1, 5, 9, and 24 hours after cells were exposed to the Nil, Cel, and ATc conditions. (**E**) Evaluation concentrations of cells that were mCherry-active in permissive and non-permissive conditions after 24 hours. Based on the mCherry fluorescence distribution of counts in the flow cytometry histograms, we determined the number of counts that was mCherry-active; representative raw data are shown in **Supplementary Figure 1**. Each cell count was divided by the sample volume used in flow cytometric analysis to determine the mCherry-active cell concentration. An asterisk (**\***) indicates that the cell concentration is lower than, or has no significant statistical difference from, the noise level from wild-type *E. coli* cells (P > 0.2 from two-tailed unpaired t-test).

We developed this strategy in *Escherichia coli Nissle 1917* (*EcN*), which is a well-studied probiotic strain that has been engineered for diagnosis and treatments of infectious diseases (*30*), gut inflammation (*31, 32*), tumors (*33, 34*), and a genetic disease (*1*). The performance of our biocontainment system was fully maintained in engineered *EcN* for at least 14 days under the permissive condition, which is the presence of a disaccharide, cellobiose; the target exogenous gene in the plasmid was eliminated upon the absence of the permissive signal. Additionally, engineered EcN with and without this biocontainment system had similar growth rates, suggesting that our system did not affect cell fitness. We also demonstrated that our system generated desirable biocontainment behavior in a mouse gastrointestinal model, in which mice without cellobiose in their drinking water lost detectable engineered activities in their fecal and cecal samples, while another group of mice supplied with cellobiose had the activities maintained. Our results support that our strategy can avoid common undesirable effects from genetic safeguard devices, including genetic instability and reduced fitness, while it is efficient in sensing an environmental change to eliminate target synthetic microbial activities. With these advantages, the engineered genes are unlikely to escape as the biocontainment system can continuously degrade its target without burdening the host cells.

## RESULTS

### Development of the genetic circuit in *E. coli K-12* for monitoring the permissive signal

Our goal is to establish a genetic circuit that allows engineered cells to maintain a target plasmid in the presence of cellobiose and destroy the plasmid when this permissive signal is absent. To evaluate the feasibility of our circuit design, we first applied it to *E. coli K-12*, which is a common lab strain of bacteria. The circuit involves a two-layered transcriptional regulation design hosted by a plasmid with an ampicillin-resistance cassette (**Figure 1A**), in which a transcriptional regulator gene, *celR*, is constitutively expressed and this regulator controls the activities of the engineered promoter *P*_*LcelO*_ (*35*) that drives the expression of another regulator gene, *tetR*. TetR is a protein regulator that represses the expression of the output by interacting with the promoter *P*_*LtetO*_ (36). As CelR responds to cellobiose as an inducer, the presence of this molecule is required for continuous TetR expression, such that TetR can repress the output expression.

To characterize and optimize the genetic behavior of this permissive signal-monitoring system, a *mcherry* fluorescent protein gene is used as the output. We then developed 8 versions of the circuit with 4 and 2 ribosomal binding sites (RBSs) for CelR and TetR, respectively; each RBS has a different strength for translational activities as characterized in previous studies (*37*). The performance of these circuit derivatives was assessed with a flow cytometric method to determine expression levels of mCherry (**Figure 1B**). Cells were exposed to no inducer (Nil), cellobiose (Cel), or anhydrotetracycline (ATc; which is the inducer of TetR). Among these 8 versions, some of them did not express the output even with the absence of cellobiose (such as the version with *BBa_B0031* for *celR* and *BBa_B0034* for *tetR*), which suggests that expression levels of CelR were too low to effectively repress TetR expression.

Conversely, some versions generated high levels of output even in the presence of cellobiose (such as the version with *BBa_B0030* for *celR* and *BBa_B0031* for *tetR*), which implies that levels of TetR in these versions were too low to repress output expression. Based on these results, the version with *BBa_B0034* for *celR* and *BBa_B0034* for *tetR* was selected for creating the biocontainment system, which generated a low mCherry expression in the presence of cellobiose and a high mCherry expression in the absence of this inducer.

### Development of the biocontainment system in *E. coli K-12*

Next, we used the selected version of the permissive signal-monitoring circuit shown in **Figures 1A** to control CRISPR system expression as the output, which contains a *cas9* and *gRNA* gene for targeting the *mcherry*-containing plasmid for cleavage (**Figure 1C**). Our goal is to evaluate the feasibility of using cellobiose as a permissive signal for maintaining a target gene in the host cells, and removing the gene when cellobiose is absent. In this system, the *cas9* and *gRNA* are each driven by an independent but identical *P*_*LtetO*_ promoter. Thus, their expression is supposed to be only repressed in the presence of cellobiose. These engineered cells also contained a *mcherry* gene in another plasmid, which was constitutively expressed; this *mcherry*-containing plasmid had a kanamycin-resistance cassette and was the target plasmid of the CRISPR system for degradation (**Figure 1C**). We first cultured these engineered cells with the presence of cellobiose; additionally, ampicillin and kanamycin were added to the culture to avoid microbial contaminations. However, when these cells were tested for biocontamination behavior, the loss of target *mcherry*-containing plasmid also led to the loss of kanamycin resistance. Therefore, when they were transferred to the permissive (with cellobiose) and non-permissive conditions (no inducers or with ATc), these three new cultures only contained ampicillin but not kanamycin. Among them, only cells exposed to cellobiose maintained high mCherry expression after 24 hours (**Figure 1D**); the fluorescence of cells exposed to no inducers or ATc dropped to levels similar to that of wild-type cells.

For cells exposed to ATc, mCherry levels dropped significantly after 5 hours, which is expected as TetR was induced to quickly express the CRISPR system to destroy the target *mcherry*-containing plasmid. For cells that were not exposed to any inducers, CelR was active for repressing TetR expression, and thus, TetR levels gradually decreased, which eventually became insufficient to repress the CRISPR system; after 9 hours in the no inducer condition, these unexposed cells reached basal levels of mCherry fluorescence and they maintained that in the culture at 24-hour. For cells exposed to cellobiose, the CelR regulator did not repress *tetR* expression, which allowed high TetR protein levels to repress the CRISPR activities, such that the *mcherry* gene was maintained in these cells.

We also assessed the levels of cells expressing the mCherry protein after exposing to permissive (Cel) and non-permissive (Nil and ATc) conditions for 24 hours (**Figure 1E**). We used flow cytometry to determine the concentrations of cells that were mCherry fluorescent in these cultures (**Supplementary Figure 1**). Our method is to collect 50,000 counts that are expected to be bacterial cells; among these counts, we determined the number of mCherry-active cells and the volume of sample used for the analysis, which were the basis for calculating the cell concentration (**Figure 1E**). With samples of cells in the permissive condition (Cel), over 10^9^ cells/mL were mCherry-active. In contrast, samples of cells in non-permissive conditions (Nil and ATc) had a signal level similar to the background noise from wild-type *E. coli* cells (Control). These results suggest that our genetic biocontainment design is feasible for constraining a target heterogeneous cellular function in a specific permissive environment.

### Development of the biocontainment system in *EcN*

After demonstrating the feasibility of our biocontainment design, we implemented it in *EcN* to demonstrate that our design can be robustly functional in this probiotic strain to eliminate target genetic materials when the environment becomes non-permissive. We first explored whether the permissive signal detection circuit design, illustrated in **Figure 1A**, can function in *EcN*. Those plasmids developed in *E. coli* were transformed into *EcN* cells to characterize eight versions of the monitoring circuit with all combinations of RBSs *BBa_B0030, BBa_B0031, BBa_B0032*, or *BBa_B0034* for driving *celR* expression and *BBa_B0034* or *BBa_B0031* for *tetR* expression; the system controlled the expression of a *mcherry* gene driven by a *P*_*LtetO*_ promoter (**Figure 2A**). Among these eight versions, the trend of expression behavior in *EcN* was highly similar to that in *E. coli*, suggesting that these genetic parts are highly transferrable between the two strains. The version with *BBa_B0034* for both *celR* and *tetR* provided the most desirable output expression behavior, generating high output expression with no inducer or with ATc; and with the presence of cellobiose, the levels of mCherry fluorescence were close to the basal signal levels in wild-type *EcN* cells. This version of monitoring circuit was then selected for constructing the biocontainment system in *EcN*, generating a two-plasmid system (same as the system shown in **Figure 1C**)—the biocontainment plasmid provided ampicillin resistance and it contained the circuit for controlling CRISPR expression in response to permissive and non-permissive conditions for the destruction of the target plasmid; the target plasmid contained the *mcherry* gene and kanamycin-resistance cassette for assessing biocontainment activities. These two plasmids developed in the *E. coli* experiments were transformed into *EcN*. (r1_5)

**Figure 2.**
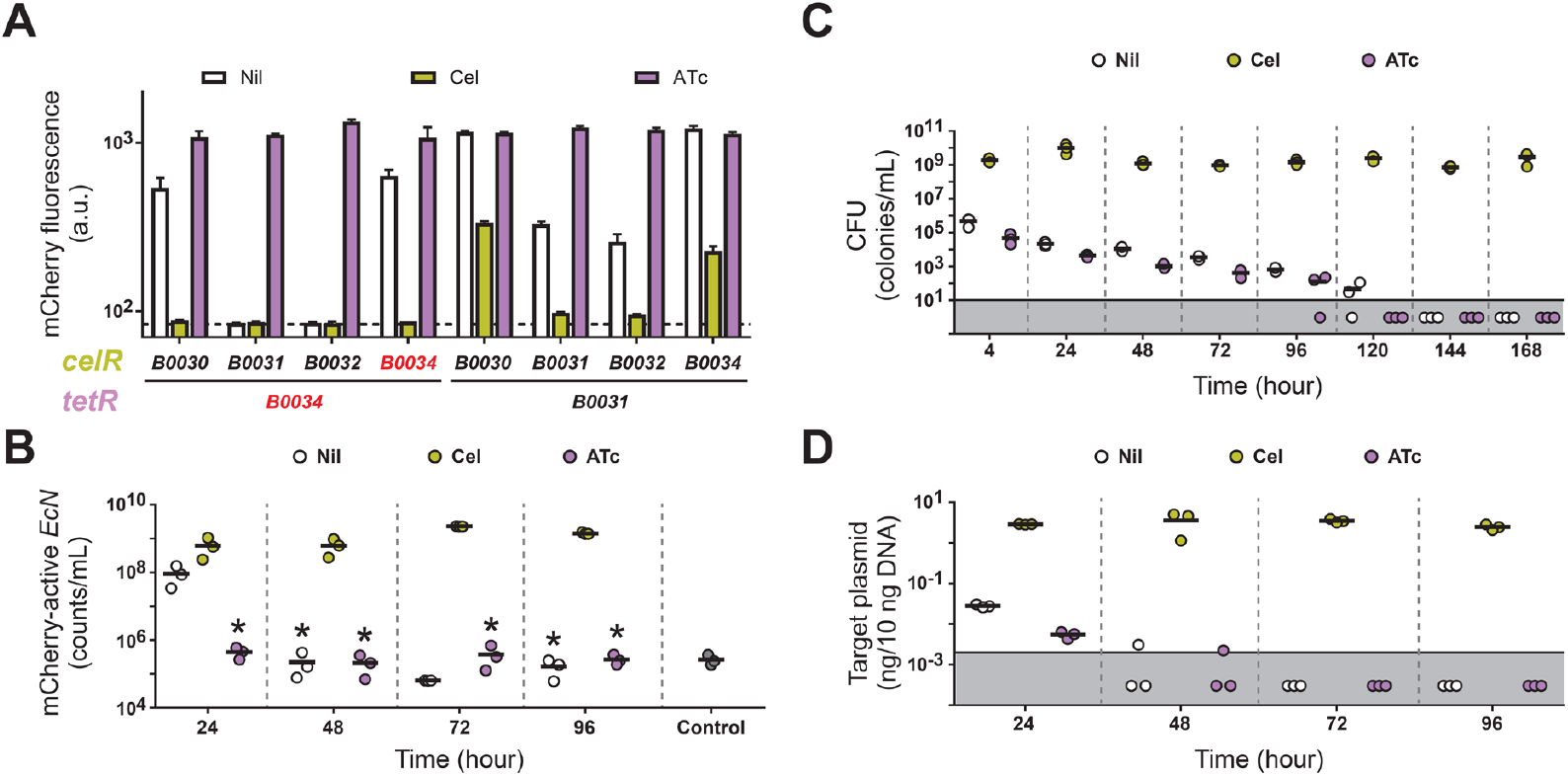
Development of biocontainment system in *EcN*. (**A**) Characterization of the permissive signal-monitoring circuit in *EcN. EcN* cells hosted the 8 versions of the circuit with different RBSs, which were the same as those illustrated in **Figure 1A**. These strains were exposed to the three conditions, no inducers (Nil), with cellobiose (Cel), and with anhydrotetracycline (ATc), and their mCherry fluorescence levels were measured with flow cytometry. Background fluorescence levels observed in wild-type *EcN* cells are shown with the dotted line. The RBSs selected for constructing the finalized circuit are colored in red, which was used to develop the two-plasmid biocontainment system in *EcN* (the plasmids are the same as those in **Figure 1C**). Each data point represents the mean ± S.D. of three biological replicates. (**B**) Characterization of biocontainment performance by monitoring mCherry expression activities. Engineered *EcN* cells were cultured in permissive (Cel) and non-permissive conditions (Nil and ATc) for 4 days and samples were collected at 24-, 48-, 72-, and 96-hour time points. The concentrations of mCherry-active cells were measured by flow cytometry. An asterisk (**\***) indicates that the flow cytometric count is lower than, or has no significant statistical difference from, the noise levels in control samples (P > 0.2 from two-tailed unpaired t-test), which were wild-type *EcN* cells with no *mcherry* gene. For **panels B to D**, each marker represents data from a biological replicate; the average of three biological replicates is illustrated with a bar. (**C**) Changes in kanamycin resistance due to biocontainment activities. To measure the loss of kanamycin resistance as a result of target plasmid degradation, colony forming unit (CFU) analysis was performed in the presence of kanamycin. The plot shows the CFU of culture samples with permissive and non-permissive conditions. For **panels C and D**, the detection limit is illustrated with the black line and a marker in the grey area below the line represents the CFU from a sample was below the detection limit. (**D**) Relative quantification of the target *mcherry*-containing plasmid in *EcN* cultures. Plasmids were isolated from cultures under those three conditions at the four time points, 24-, 48-, 72-, and 96-hr. An amount of 10 ng of each isolated plasmid sample was used to perform RT-PCR to determine the quantity of the target plasmid relative to the total plasmid in these cells.

We first characterized the sensitivity of our *EcN* biocontainment system to cellobiose (**Supplementary Figure 2**), investigating the range of cellobiose concentrations that can efficiently maintain the target plasmid. We first grew the engineered cells in culture media with 5 g/L cellobiose, ampicillin, and kanamycin; the saturated cultures were diluted 200 folds in culture media with ampicillin (no kanamycin) and a range of cellobiose concentrations to grow for 48 hours with a 1:200 dilution at 24 hours. Expression of mCherry in these cultures was assessed at 24 and 48 hours by a flow cytometric method (see Methods section). The percentage of cells expressing mCherry decreased dose-dependently in the range of 0.01 to 1.25 g/L of cellobiose. The drop in the mCherry-active portion was more significant at the 48-hour time point. When cellobiose concentration was above 2.5 g/L, there was no significant statistical difference in the resulting populations, and above 95% of cells were mCherry-active. We decided to use 5 g/L cellobiose as the permissive condition in all experiments.

Next, we tested whether *EcN* cells maintained the target *mcherry*-containing plasmid without kanamycin. As our engineered *EcN* contained two plasmids (**Figure 1C**), we normally cultured them with both ampicillin and kanamycin, preventing microbial contamination and ensuring all cells contained both plasmids. However, during the biocontainment activities, cells in non-permissive conditions are expected to lose the target plasmid and thus, also lose kanamycin resistance. Therefore, when cells were transferred to permissive and non-permissive conditions, we did not provide kanamycin to avoid cell killing. This design led us to ask whether the absence of kanamycin would promote the loss of our target plasmid due to the loss of selective pressure. Generally, many microbes maintain non-essential plasmids for an extended period even without selective pressure, and removing plasmids requires various plasmid curing methods (*38*) (r1_4). Also, *EcN* keeps non-essential plasmids such as pMUT1 and pMUT2 (*39*). To evaluate the stability of our target plasmid, we grew *EcN* containing the plasmid in our standard in vitro conditions for five days without kanamycin; cells were passaged (diluted 200 folds) once each day, and the mCherry levels of overnight cultures were assessed (**Supplementary Figure 3**). Our flow cytometric analysis shows that the *EcN* population consistently had 90 to 98% with mCherry expression over the 5 days, supporting that *EcN* maintained our target plasmid even without kanamycin.

### Assessment on biocontainment efficiency

Based on these validations, we designed the experiments to evaluate the efficiency of our *EcN* biocontainment system. Engineered *EcN* cells were cultured overnight in media with 5 g/L cellobiose, ampicillin, and kanamycin. The saturated cultures were diluted (1:200) into three conditions that had ampicillin but no kanamycin for performing biocontainment activities, which included the permissive condition (with cellobiose) and two non-permissive conditions (no inducers or with ATc). Each culture was grown for multiple days in the same condition with a passaging (1:200 dilution) once each day. In each day, samples were collected from these cultures to analyze the efficiency in destroying the target plasmid. Three methods were used, including flow cytometry to determine mCherry expression, colony forming unit (CFU) analysis to measure the concentration of cells that remained kanamycin-resistant, and real-time quantitative PCR (RT-PCR) to quantify the amount of *mcherry* gene DNA fragments in cells.

For flow cytometric analysis, we collected samples from saturated cultures each day, ensuring cells had sufficient time to express and accumulate mCherry protein. We acquired 50,000 events that were expected to be from *EcN* cells based on side scattering and forward scattering properties. The software then determined the number of counts that were mCherry-active based on fluorescence intensity. With the information of sample volume analyzed and the dilution factor, we calculated the concentration of mCherry-active *EcN* cells in the culture. To evaluate the limit of quantification, we first used this method to analyze wild-type *EcN* cells as control samples, which did not contain any *mcherry* genes (**Figure 2B**). In three biological replicates, the background noise was 3, 4, and 6 counts within the 50,000 collected events (about 0.03%), which is equivalent to (2.6 ± 0.9) ×10^5^ counts/mL; this background noise can be due to random photon emission and detection (*40*).

We then used this flow cytometric method to assess changes in mCherry expression activities in response to permissive and non-permissive conditions (**Figure 2B**). For cultures with cellobiose, which is the permissive condition, the concentration of mCherry-active *EcN* cells maintained around 109 cells/mL throughout the 4-day experiment. In contrast, those cultures exposed to ATc, which directly induced TetR regulator for CRISPR expression, reached the background level (about 10^5^ counts/mL) at the first measured time point (24-hour) and the concentration of mCherry-active signals remained at this level in subsequent days. With no inducers, which is also a non-permissive condition, mCherry-active population decreased about 10 folds (10^8^ cells/mL) after 24 hours; it then reached the background level at 48-hour and remained at that level in all subsequent time points. The flow cytometric method has the advantage of direct measuring protein expression levels of the target engineered gene; however, due to the relatively high level of background noise, its sensitivity is also relatively low. We then explored additional methods to assess the biocontainment efficiency.

To further evaluate the biocontainment performance, we used colony-forming unit (CFU) analysis to determine the *EcN* population with kanamycin resistance (**Figure 2C**). In addition to a *mcherry* gene, the target plasmid for degradation also contained a kanamycin resistance cassette and thus, the biocontainment activities can be assessed by monitoring this antibiotic resistance within the population. To ensure cells were active, each day, we diluted each overnight culture 200 folds in the same condition (Nil, Cel, or ATc); as these cultures did not contain kanamycin, cells that already lost the target plasmid still grew normally. However, when these fresh cultures reached an optical density at 600 nm (OD600) of about 1, they were serially diluted and then spotted on LB agar solid media with kanamycin, cellobiose, and ampicillin, where engineered *EcN* cells could form colonies if the target plasmid was not degraded. The highest number of cells used in this CFU analysis was by using 100 µL of undiluted culture, which provided a limit of quantification at 10 colonies/mL.

Results from this CFU method show that engineered *EcN* lost kanamycin resistance only in non-permissive conditions but not in the permissive condition (**Figure 2C**). For cultures with cellobiose (the permissive condition), CFU was above 10^9^ cells/mL at all time points, suggesting the target plasmid was maintained in the population. For biocontainment activities, after the original cultures were diluted in non-permissive conditions and grew to an OD600 of about 1 (which took about 4 hours), cultures with no inducers had a CFU dropped to (4.7 ± 2.3) ×10^5^ colonies/mL (4-hour time point on **Figure 2C**); that of ATc samples was below 10^5^ colonies/mL.

For many biocontainment systems that kill equipped cells, they generate cellular stress that promote pathways for mutating the genetic device; thus, cells that survived from the initial killing phase usually have silenced the biocontainment function, becoming escapees to dominate the microbial population. In contrast, our mechanism is not expected to hamper any endogenous pathways, which favors the biocontainment system to continually scavenge the remaining target plasmid for destruction. Indeed, CFU of non-permissive cultures continued to decrease throughout this experiment while that of permissive cultures (Cel) maintained above 10^9^ colonies/mL, supporting that the biocontainment system kept functional for multiple days (**Figure 2C**). After 144 hours in both non-permissive conditions, no kanamycin-resistant *EcN* cells were detected in all biological replicates. These results suggest that destroying engineered genes, instead of killing engineered cells, can be a promising strategy for avoiding escapee formation.

In addition to assessing engineered cellular functions, we also used real-time PCR (RT-PCR) to quantify the amount of *mcherry* DNA fragments in these cultures (**Figure 2D**). Samples were collected each day from saturated culture, such that the target *mcherry*-containing plasmid were accumulated to its highest cellular level. Plasmids were isolated from each sample, which could contain the biocontainment plasmid and the target *mcherry*-containing plasmid (**Figure 1C**). An amount of 10 ng of each total plasmid sample was used to perform RT-PCR with a pair of primers that targeted the *mcherry* gene. Our standard curve (**Supplementary Figure 4**) demonstrates that this method robustly quantified the *mcherry*-containing plasmid with a range of 2 pg to 6 ng of the target plasmid (per 10 ng of total plasmid). However, when the quantity of the plasmid was 1 pg or below, the C_t_ values varied significantly among technical replicates; thus, we expect the quantification limit to be close to 2 pg in 10 ng of plasmid DNA.

With RT-PCR, we showed that the target plasmid was degraded under non-permissive conditions (**Figure 2D**). With the permissive condition, the quantity of the target plasmid was above 1 ng in 10 ng of isolated DNA at all time points. For engineered *EcN* in non-permissive conditions, after 24 hours, target plasmid levels were reduced to (2.7±0.2) ×10^−2^ ng and (5±1) ×10^−3^ ng per 10 ng DNA in Nil and ATc samples, respectively. After 72 hours, based on measured C_t_ values, the quantity of the target plasmid was below the quantification limit (2 pg/10 ng isolated DNA) in all biological replicates under both non-permissive conditions.

Together, the results from flow cytometry, CFU analysis, and RT-PCR support that the biocontainment system efficiently destroyed the target plasmid in non-permissive conditions but not in permissive conditions, which ceased the engineered biological activities. This biocontainment strategy provides a unique advantage in preventing escapee formation as it enables continuous elimination of the target for multiple days.

### Assessment of cell fitness

After characterizing the efficiency of the biocontainment system, we evaluated the influence of this system on cellular fitness (**Figure 3**). For many genetic safeguard systems, their mechanisms involve killing engineered cells; the basal activities of these safeguards are often harmful to the host cells, dampening cell fitness. Also, they generate cellular stress that promotes genetic instability to mutate and deactivate these biocontainment systems, leading to short lifetime of the genetic device and high levels of escapees (*20, 21*). We hypothesize that our biocontainment strategy does not have this weakness as it is not expected to interfere with endogenous cellular activities to create stress. We evaluated this aspect by analyzing growth rates (**Figure 3A**); additionally, we tested the stability of the system with a 14-day experiment (**Figures 3B and 3C**).

**Figure 3.**
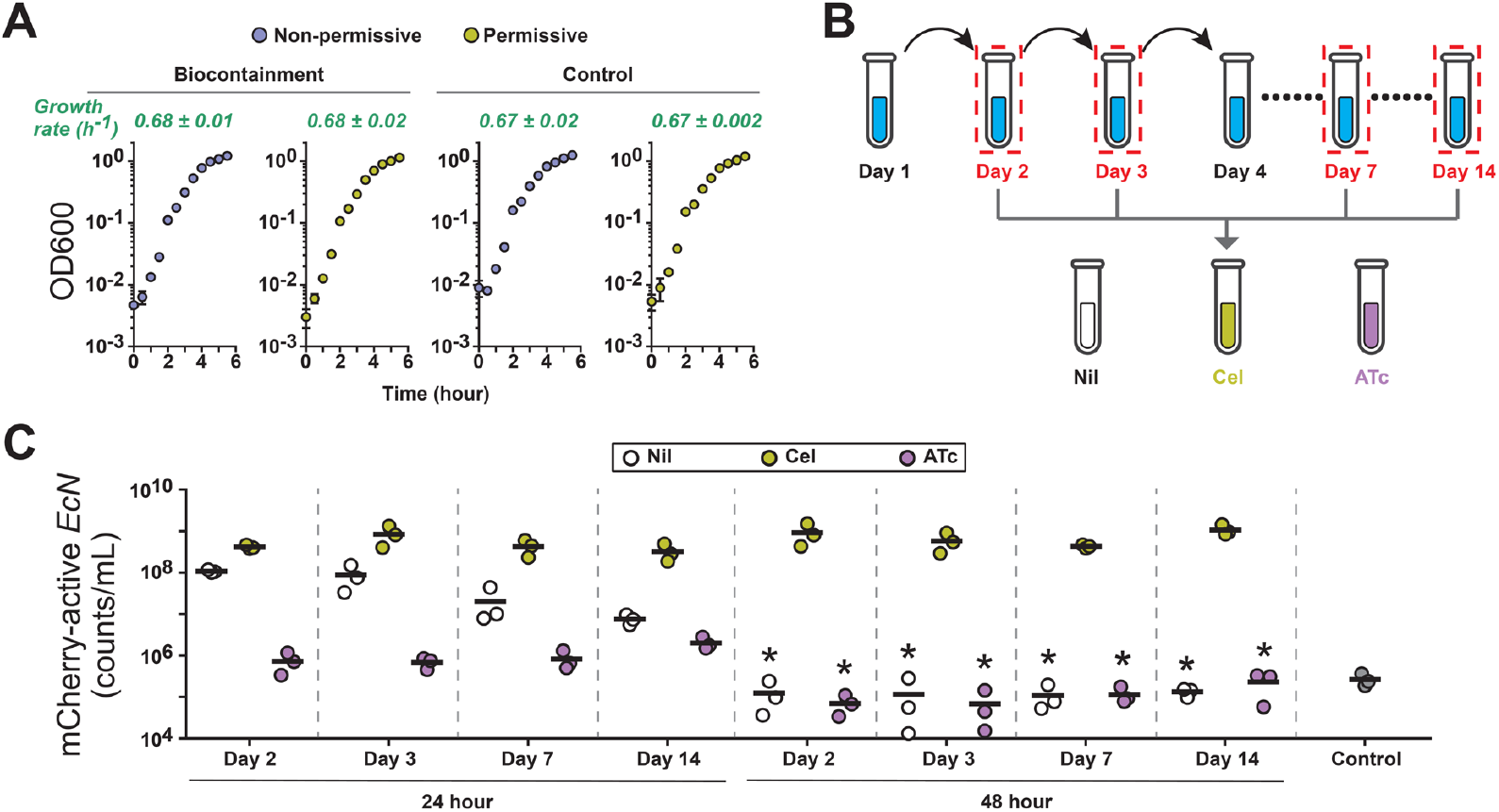
Cell fitness of engineered *EcN*. (**A**) Growth rates of two engineered *EcN* strains. The Biocontainment strain contained both the Biocontainment plasmid and the Target plasmid as shown in **Figure 1C**; the Control strain only contained the Target plasmid. The two strains were grown without any inducer (non-permissive) and with cellobiose (permissive). Each data point represents mean ± S.D. of OD600 measurements from three biological replicates; the error bar is not shown if it is shorter than the marker. The growth rate of each culturing condition (mean ± S.D. in green color) was calculated with data obtained at 1 and 3 hours. (**B**) Schematic diagram of the 14-day experiment for characterizing the biocontainment system in *EcN*. Engineered cells were grown and passaged for 14 days and they were used to characterize the biocontainment function on Days 2, 3, 7, and 14 by exposing to permissive (Cel) and non-permissive (Nil and ATc) conditions for 48 hours, with a passaging at 24 hours. Samples were collected at 24 and 48 hours for flow cytometric analysis. (**C**) Characterization of mCherry expression activities from engineered *EcN*. After exposing cells to the three conditions for 24 and 48 hours, their levels of mCherry fluorescence were measured with flow cytometry. In the plot, each marker represents the cell concentration that was active in mCherry expression in a biological replicate. The average of three biological replicates is shown as a bar. An asterisk (*) indicates that the flow cytometric count is lower than, or has no significant statistical difference from, the noise levels in control samples (P > 0.2 from two-tailed unpaired t-test), which were wild-type *EcN* cells with no *mcherry* gene.

We compared the growth rates of engineered *EcN* with and without the biocontainment plasmid at permissive and non-permissive conditions (**Figure 3A**). We prepared overnight cultures of *EcN* hosting either both the biocontainment plasmid and target plasmid (Biocontainment strain), or only the target plasmid (Control strain); both strains were grown in the presence of cellobiose but Biocontainment strain was grown in the presence of ampicillin and kanamycin and the Control strain in only kanamycin. The use of antibiotics ensures cells contained those plasmids at the beginning of growth rate analysis. These overnight cultures were diluted 200 folds to grow in the presence of cellobiose as the permissive condition or with no inducer as the non-permissive conditions. To ensure cells continue to maintain the biocontainment system, the Biocontainment strain cultures still contained ampicillin but not kanamycin; the Control strain cultures had no antibiotics. Optical density at 600 nm (OD600) was measured for 5.5 hours to assess cell growth. The growth activities of all these cultures were similar, where OD600 increased exponentially between the 1-to 3-hour time points; after that, cell growth slowed down to enter the stationary phase. Exponential growth rates in all four conditions were 0.67 to 0.68 h^-1^ without significant difference between the two strains based on two-tailed unpaired t-test analysis (P > 0.2). These results support that our biocontainment system did not affect cell growth when both activated or not activated.

Additionally, the stability of the biocontainment system was evaluated with a 14-day experiment (**Figure 3B**). Cells were cultured at the permissive condition, with the presence of ampicillin and kanamycin, and passaged once per day for 14 days. On Days 2, 3, 7, and 14, cells were diluted 200 folds into the two non-permissive conditions, including no inducer and with ATc; as control experiments, each culture was also diluted into the permissive condition (with cellobiose). These cultures only contain ampicillin but no kanamycin. These cultures were again diluted once each day and after 24 and 48 hours in these conditions, we used the flow cytometric method to assess mCherry expression activities as described in **Figure 2B**. Corresponding results suggest that the biocontainment performance was consistent throughout the 14 days (**Figure 3C**)—engineered *EcN* in non-permissive conditions showed significant decreases in the mCherry-active population after 24 hours and at the 48-hour time point, the mCherry-active population dropped to background noise levels (this background was determined from wild-type *EcN* cultures with no *mcherry* gene). Representative raw data from Day 14 assessment were illustrated in **Supplementary Figure 5**. Cellular responses to non-permissive conditions were highly similar on all the assessed days, suggesting the biocontainment system was stable and robust for at least 14 days.

### Assessment of biocontainment efficiency in a mouse gastrointestinal environment

After in vitro characterization of the *EcN* biocontainment system, we explored its capability in a mouse gastrointestinal model as described in **Figure 4A**. For synthetic probiotics that have therapeutic functions, it is desirable that the engineered activities can be quenched in patients either after or during therapy. To evaluate our biocontainment system for this application, we assessed its use for maintaining engineered genetic activities in a mouse gut environment with a continuous dietary supply of the permissive signal, cellobiose, and for removing the gene to stop the activities by taking away the signaling molecule from the diet.

**Figure 4.**
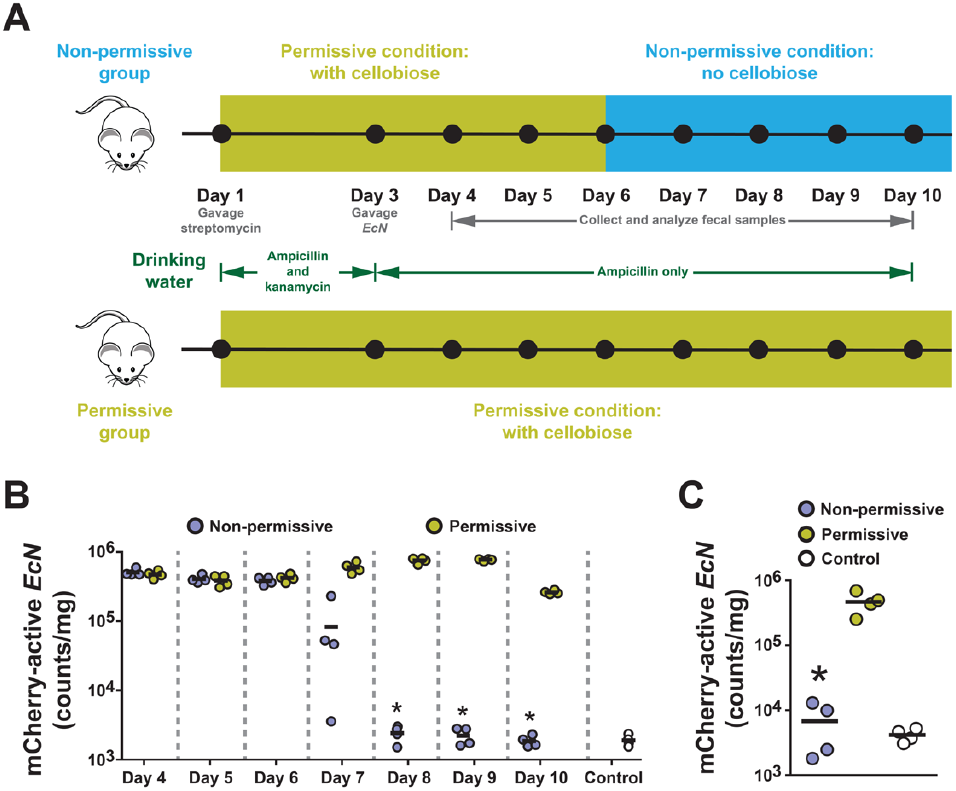
Characterization of engineered EcN in a mouse gastrointestinal environment. **(A)** A schematic diagram of animal studies. To enhance *EcN* colonization, three antibiotics were provided to mice, including streptomycin by oral gavage on Day 1, ampicillin and kanamycin via drinking water ad libitum from Day 1 to 3, and ampicillin only via drinking water ad libitum from Day 4 to 10. An amount of 5*10^8^ CFU of *EcN* equipped with the biocontainment system was orally gavaged to mice on Day 3. Fecal samples were collected once per day on Days 4 to 10 for assessing mCherry expression and kanamycin-resistant activities in the microbiome. At the end of the experiments, cecal samples were collected after sacrificing these mice. For the non-permissive group (n = 4), the permissive signal, cellobiose, was only provided on Days 1 to 6 via drinking water. For the permissive group (n = 4), cellobiose was provided throughout the experiments, from Days 1 to 10. **(B)** Flow cytometric analysis of fecal samples. The plot illustrates the count per mg of fecal sample that are considered mCherry-active; representative histograms of mCherry fluorescence distribution are presented in **Supplementary Figure 4**. For panels **B** and **C**, each data point is illustrated with a marker; the average of 4 biological replicates is shown as a bar. An asterisk (*) indicates that the cell concentration is lower than, or has no significant statistical difference from, the noise levels in control samples (P > 0.2 from two-tailed unpaired t-test), which were isolated from mice not treated with EcN. **(C)** Expression activities of *mcherry* in cecal samples. Flow cytometric analysis was also performed on cecal samples to determine the *mCherry* expression in the gut microbiomes.

To create a permissive environment in the gut, mice were continuously provided with drinking water that contained cellobiose starting on Day 1 of the experiment. In parallel, we provided 20 mg of streptomycin to mice by oral gavage, an antibiotic that is expected to reduce the microbiome population, enhancing the colonization of our engineered *EcN* (*19*). Additionally, in Days 1 to 3, ampicillin and kanamycin were provided via drinking water, aiming to continually reduce native microbiome population; these two antibiotics were selected as our engineered *EcN* cells were initially resistant to them; however, cells are expected to lose the target plasmid and kanamycin resistance when they were in non-permissive conditions. Therefore, after gavaging the engineered cell to these mice on Day 3, we ceased the supply of kanamycin in drinking water, ensuring the gut would not contain this antibiotic when it is switched to the non-permissive condition; *EcN* should keep the target plasmid even in the absence of kanamycin, as we demonstrated in **Supplementary Figure 3**. Ampicillin supply was continued to facilitate the colonization of engineered *EcN* in the gut.

After the gavage of *EcN* cells, fecal samples were collected once per day to evaluate mCherry expression activities in the microbiome. In Days 4 to 6, cellobiose was continuously supplied to all mice via drinking water and high mCherry levels were consistently observed in their fecal samples (**Figure 4B**), supporting that engineered *EcN* was inhabiting in the gut. After collecting fecal samples on Day 6, these mice were split into a non-permissive group (n=4), where cellobiose was not supplemented in drinking water, and a permissive group (n=4) with cellobiose provided as before. To determine the background noise levels in flow cytometric measurements, fecal samples were also collected and analyzed on Day 10 from a group (n=4) of mice that were not treated with *EcN*. Bacterial cells isolated from these untreated mice were from the native microbiomes; it is expected that native bacteria were present in all mouse samples, contributing to a majority of cell counts with no mCherry expression. The data served as controls to determine the limit of quantification, which was (1.8 ± 0.3) ×10^3^ counts/mg fecal sample.

For the non-permissive group, levels of mCherry-active cells dropped significantly on Day 7 and they were not detectable since Day 8 based on the levels of background noise from control samples (**Figure 4B**). In contrast, all samples from the permissive group contained mCherry-active cells throughout this experiment.

At the end of the experiment, we analyzed cecal samples (**Figure 4C**) and the results are consistent with the fecal analyses, where samples from the permissive group contained mCherry-active cells but not those from the non-permissive group. These mice studies strongly support that our biocontainment is efficient in a gastrointestinal environment—a supply of cellobiose in the diet maintained the engineered cellular function and that function was terminated by discontinuing the cellobiose supply. Representative raw data from flow cytometric analysis of fecal and cecal samples are illustrated in **Supplementary Figure 6**.

In addition to flow cytometric analysis, we used CFU analysis to measure the levels of cells with kanamycin resistance in fecal and cecal samples (**Supplementary Figure 7**), further evaluating biocontainment activities in the mouse gastrointestinal model.

Similar to the approach in the in vitro studies (**Figure 2C**), we serially diluted bacterial samples isolated from feces and cecum contents and then spotted these dilutions onto solid culture media with kanamycin, ampicillin, and cellobiose for colonies to form. To determine the quantification limit, we analyzed control samples from mice without *EcN* treatment; some colonies were formed from these control samples, suggesting that the native microbiomes contained microbes with resistance to kanamycin and ampicillin. The levels of these native microbes in fecal and cecal samples were (1.3 ± 1.6) ×10^3^ and (1.7 ± 1.4) ×10^4^ colonies/mg samples, respectively (**Supplementary Figure 7**). Results from CFU analysis complement the flow cytometry data in **Figure 4B**, in which fecal samples from the permissive group maintained a CFU of around 10^5^ to 10^6^ colonies/mg throughout the experiment; in contrast, the CFUs of fecal samples from the non-permissive group dropped to the background level after cellobiose was not supplied (Days 8 to 10). Similar pattern was observed from the CFUs of cecal samples. These results further confirm that the biocontainment system is functional in a gastrointestinal environment.

## DISCUSSION

It is necessary to gain reliable control of engineered activities in synthetic probiotics to ensure host safety and avoid their transmission to other hosts. Conventionally, researchers aim to achieve this goal by implementing genetic kill switches to kill engineered cells or suppress their growth under non-permissive conditions; basal activities of these systems may constitutively generate cellular stresses, which hamper cellular fitness and promote genetic instability. There is a strong selective pressure to mutate and silence these kill switches, leading to a high frequency of escapees.

The biocontainment approach that we created addresses this problem by specifically targeting the synthetic gene for removal, instead of destroying any endogenous cellular components. This approach promotes the orthogonality between the biocontainment activities and the rest of cellular functions, which minimizes the burden on engineered cells. As the destruction of target DNA does not obstruct endogenous cellular activities, the biocontainment system is stable and can perform continuously for an extended period until the target is eliminated. This is demonstrated in **Figure 2C**, in which our biocontainment system kept reducing the kanamycin-resistant population over multiple days and finally became undetectable.

Our strategy depends on the specific removal of DNA fragments in engineered cells, which can be achieved by various genetic tools. For example, DNA recombinase systems, such as Cre-loxP, excises target genes in between two loxP sites (*41*). However, these tools recognize specific DNA sequences and thus, they require the incorporation of specific elements in the target for DNA excision. To avoid the need to modify biocontainment targets, we selected CRISPR system; sequence-specific DNA destruction can be robustly attained by expressing a tailored guide RNA (gRNA) to recognize and cleave a target genetic material. Previous studies demonstrated CRISPR can efficiently remove target DNA sequence in microbes; for instance, Caliando et al. used an inducible transcriptional expression system to control a CRISPR tool; upon the presence of the inducer for CRISPR expression, target plasmid DNA was degraded (*28*). Hayashi et al. showed that a constitutively expressed CRISPR system can be used as a tool to prevent their engineered bacteria from gaining a specific gene via horizontal gene transfer (*29*). Differing from these studies, we aim to activate the CRISPR activities whenever a specific signal is absent, such that the target gene can be maintained in engineered cells only under a permissive condition defined by that signal.

For sensing and responding to permissive and non-permissive conditions, we constructed a two-layered transcriptional regulatory circuit by using allosterically regulated transcriptional repressors (**Figure 1A**). This design has been employed in several kill switches to implement a Boolean NOT function (*16, 19, 42*), in which an expression of the cytotoxic output only occurs in the lack of the input. A key advantage of the circuit-based approach is the high modularity of the output; instead of controlling toxic gene expression, we harnessed this circuit design to control CRISPR activities for degrading the target plasmid (**Figure 1C**).

Our circuit was developed with the transcriptional repressor CelR so that cellobiose is the permissive signal. Cellobiose was selected as it is unlikely to lead to any adverse effects on the human gastrointestinal system. It is a disaccharide indigestible by human enzymes but can be metabolized by a spectrum of gut microbes in the colon. Previous studies show that high-dose oral administrations of cellobiose are safe for human beings (*43, 44*) and may increase the population of beneficial microbes in the microbiomes that contribute to gut health, such as *Lactobacillaceae* species (*45*). These properties render cellobiose as an appropriate permissive signal for controlling engineered *EcN* in the gut environment. Indeed, our results suggest that cellobiose has great potential to serve as a signaling molecule for synthetic microbes in gastrointestinal tracts. However, it is highly possible that cellobiose may not reach some parts of the host system, which limits its uses. For example, Din et al. (*46*) engineered microbes that can be orally administered to inhabit in the liver for drug delivery; it is unclear on the accessibility of cellobiose to these disease sites for biocontainment. Therefore, further studies are required to determine the feasibility of our permissive condition design for each specific application.

To serve as proof of this concept, we used our design to target the removal of a plasmid that contains a kanamycin resistance cassette and a *mcherry* gene. Our in vitro and in vivo experiments imply that our genetic method is feasible for controlling target genetic materials in the probiotics in response to an environmental change—in the gut environment, mCherry expression became undetectable in fecal samples by our flow cytometry method after 48 hours without a supply of cellobiose. Analyses of cecal samples further confirmed that the engineered activities were eliminated in the gut microbiome. In comparison, for mice with a supply of cellobiose, mCherry-active cells were consistently detected throughout the experiments after *EcN* administration. These results support that our strategy has the potential for synthetic probiotics applications.

As a main limitation, our biocontainment strategy requires additional genetic components to prevent the spread of a target plasmid by horizontal gene transfer. Our design constrains engineered cellular activities by destructing the target non-essential plasmid. However, the plasmid can potentially be transferred horizontally to other microbes in the microbiome, escaping from our biocontainment system. This problem may be reduced by including the biocontainment system and target genes in the same plasmid, such that the target plasmid can still be destroyed by the CRISPR in the recipient cells. Alternatively, previous studies have created a range of safeguard methods to avoid horizontal gene transfer. For instance, toxin-antitoxin pairs were used in previous designs, where toxin genes were incorporated into the plasmid and corresponding antitoxins were expressed in the engineered microbe. Horizontal transfer of this plasmid is prevented as recipient cells without the corresponding antitoxins are killed by those toxins (*47*).

Additionally, our genetic design involves two plasmids, with one plasmid containing the biocontainment system and the other plasmid containing the target engineered gene. The advantage of this design is the ease and flexibility of implementing the biocontainment system to different engineered cells—after replacing the gRNA of the CRISPR, the plasmid is ready to be incorporated into the engineered cells for constraining a new target. However, with our current design, the biocontainment plasmid is not targeted for destruction and it may linger in engineered cells. This risk can be eliminated by implementing an additional gRNA for removing the biocontainment plasmid. As next steps for developing this biocontainment strategy to enhance biosafety, we plan to assess its performance in a range of probiotics hosts and explore its use for targeting therapeutic genes.

## ACKNOWLEDGMENTS

This work was supported by grants from the US National Institutes of Health (NIH), including R35GM142421 by National Institute of General Medical Sciences, R41CA275454 by National Cancer Institute, and R16NS131108 by National Institute of Neurological Disorders and Stroke.

## AUTHOR CONTRIBUTIONS

N.N. and C.T.Y.C. designed and conducted microbial experiments; M.W. conducted mouse experiments; M.W. and L.L. designed mouse experiments; C.T.Y.C. wrote the paper.

## SUPPLEMENTARY INFORMATION

**Supplementary Document**. Supplementary Figures 1 to 7; Supplementary Table 1

**Supplementary Data 1**. Map and sequence of the biocontainment plasmid in the GenBank format.

**Supplementary Data 2**. Map and sequence of the *mcherry*-containing plasmid in the GenBank format.

## MATERIALS AND METHODS

### Strains of bacteria

Cloning was performed with commercial cloning strains of *E. coli*, including *ig10β* (Intact Genomics, Inc.; St. Louis, MO) and *XL-1 Blue* (Agilent Technologies, Inc.; Santa Clara, CA). Experiments were performed with *E. coli K-12 MG1655* (ATCC; Manassas, VA) and *Nissle 1917* (from Dr. James J. Collins lab at Massachusetts Institute of Technology).

### Plasmid construction

Standard cloning methods were used with primers and single-stranded DNA oligomers purchased from Eurofins Genomics LLC (Louisville, KY) and cloning enzymes and reagents from New England Biolabs (Ipswich, MA). DNA oligomers used in this study were listed in **Supplementary Table 1**. For building the biocontainment plasmid, *pZA1* (Expressys, Germany) was used as the vector backbone, which contained a *p15A* origin of replication and an ampicillin-resistant cassette. The biocontainment circuit fragment was synthesized by VectorBuilder Inc. (Chicago, IL) to incorporate into the vector, which contained the *celR, tetR, cas9, gRNA*, and the genetic elements to control the transcription of these genes; the sequence of the optimized version of this plasmid is shown in **Supplementary Data 1**. For the permissive signal-monitoring circuit, *cas9* gene fragment was replaced with a *mcherry* gene fragment from PCR via restriction sites, SphI and HindIII. To alter ribosome binding sites (RBSs), two complementing single-stranded DNA were mixed to reach a final concentration of 50 µM; the mixture was heated to 100 °C and then allowed to cool down under room temperature for hybridization. The double-stranded DNA possessed sticky 5’- and 3’-ends that mimic digested restriction sites. These RBS fragments were incorporated into the plasmid, via restriction sites BsrGI and AvrII for *celR*, and NheI and NcoI for *tetR*.

To build the *mcherry*-containing plasmid as the target of elimination, the plasmid pTR (*35*) from our previous studies was used as the vector, which contains a *ColE1* origin of replication and a kanamycin-resistance cassette. A constitutive promoter, *P*_*L*_, was cloned into this plasmid via restriction sites SalI and EagI. This promoter drove the expression of the *mcherry* gene that was cloned via EagI and BsrGI. The sequence of this *mcherry*-containing target plasmid is shown in **Supplementary Data 2**.

### Cultural conditions

All culturing reagents were purchased from VWR (Radnor, PA) and Fisher Scientific (Waltham, MA). Both *E. coli* and *EcN* were grown in LB broth (3 to 5 mL for each culture) at 37 °C and 200 rpm. Unless specified, the concentrations of reagents in cell cultures were 100 µg/mL for ampicillin, 50 µg/mL for kanamycin, 5 g/mL for cellobiose (Cel), and 50 ng/mL for anhydrotetracycline (ATc).

### In vitro characterization of *E. coli* and *EcN* biocontainment systems

For characterizing the permissive signal-monitoring system, both *E. coli* (**Figure 1B**) and *EcN* (**Figure 2A**), containing the Monitoring plasmid (**Figure 1A**), were grown overnight in LB media with ampicillin and cellobiose. The overnight cultures were diluted 200 folds with fresh LB media with ampicillin and either no inducers (Nil), Cel, or ATc (three different conditions). *E. coli* and *EcN* cells were collected after 6 and 24 hours, respectively. These culture samples were immediately analyzed with the flow cytometry method (see the Flow cytometry section below).

For the characterization of the biocontainment system in *E. coli, E. coli* cells containing the two plasmids in **Figure 1C** were grown overnight with kanamycin, ampicillin, and cellobiose.

Overnight cultures were then diluted 200 folds with the culture media containing only ampicillin (no kanamycin) with either no inducers (Nil), Cel, or ATc. Samples were collected at 1-, 5-, 9-, and 24-hour for flow cytometry.

To determine cellobiose sensitivity of engineered *EcN* (**Supplementary Figure 2**), cells with the two plasmids in **Figure 1C** were grown overnight. The overnight cultures were diluted 200 folds with culture media containing ampicillin (no kanamycin) and a series of concentrations of cellobiose, including 10, 5, 2.5, 1.25, 1, 0.63, 0.5, 0.31, 0.1, 0.05, 0.025, 0.01, and 0 g/L. Samples were collected after growing in these conditions for 24 hours; each culture was then passaged (diluted 200 folds again with the same culture condition) and samples were collected again from these new cultures after another 24 hours (48-hour time point). Collected samples were analyzed with flow cytometry.

For evaluating the stability of the target *mcherry*-containing plasmid in *EcN* without kanamycin (**Supplementary Figure 3**), colonies of *EcN* cells containing only the target plasmid (without the biocontainment plasmid) were picked from a LB agar plate with kanamycin and grown overnight with no antibiotics. In the next 5 days, these cultures were passaged in LB media with no antibiotics once each day and samples were collected from each overnight culture for flow cytometric analysis.

To assess the *EcN* biocontainment efficiency (**Figures 2B to 2D**), colonies of cells containing both the biocontainment and target plasmids (see **Figure 1C**) were cultured in LB media with kanamycin, ampicillin, and cellobiose. Overnight cultures were diluted 200 folds in media with ampicillin (no kanamycin); chemical reagents were added to these cultures to generate three conditions, including the permissive condition (with cellobiose, Cel) and two non-permissive conditions (with no inducer, Nil, or with anhydrotetracycline, ATc). For flow cytometric analysis (**Figure 2B**) and RT-PCR (**Figure 2D**), these cultures were grown for 4 day. Each day, samples were collected from overnight cultures and diluted 200 folds into fresh media with the same conditions; these diluted cultures were used the next day to repeat this process. For CFU analysis (**Figure 2C**), cultures were grown for 6 days. On each day, overnight cultures were diluted 200 folds and then grown until OD600 reached about 1 for sample collection; after collection, each remaining culture continued to grow for the next-day sample collection. Each collected sample was serially diluted for 10^0^, 10^1^, 10^2^, 10^3^, 10^4^, 10^5^, 10^6^, 10^7^ folds with PBS; 5 µL of each diluted sample was spotted on a LB agar plate with ampicillin, kanamycin, and cellobiose. Additionally, 100 µL of undiluted sample from each culture was plated, which allowed us to reach a quantification limit of 10 colonies/mL. Colony formation was recorded the next day.

To study the biocontainment stability (**Figures 3B and 3C**), engineered *EcN* cells containing both the biocontainment plasmid and target plasmid (**Figure 1C**) were grown in the presence of ampicillin, kanamycin, and cellobiose for 14 days with passaging once per day by a dilution of 200 folds. On each day, overnight cultures were also 1:200 diluted into three conditions for testing the biocontainment activities. They were transferred to culture media with only ampicillin, and these media contained either no inducer, cellobiose, or ATc. After 24 hours, these cultures were characterized with the flow cytometric method and they were passaged with a 200-fold dilution in the same conditions. At 48 hours upon transferring cells to the three conditions, the cultures were analyzed again.

### Characterization of *EcN* growth

To evaluate the influence of the biocontainment system on cell growth (**Figure 3A**), we used two *EcN* strains. One strain contained both the biocontainment plasmid and the target plasmid (biocontainment strain) and the other strain contained only the target plasmid (control strain). The biocontainment strain was first inoculated into cultural media with ampicillin, kanamycin, and cellobiose, ensuring all cells in the cultures contained both plasmids; the control strain was inoculated into media with only kanamycin and cellobiose, as it did no possess ampicillin resistance. Overnight cultures of both biocontainment and control strains were diluted 200 folds in fresh media at the permissive (with cellobiose) or a non-permissive (no inducer) condition, each new culture was 5 mL and only contained ampicillin but not kanamycin, as cells in non-permissive condition lost kanamycin resistance after the target plasmid was removed. Optical density at 600 nm (OD600) was measured at 0, 0.5, 1, 1.5, 2, 2.5, 3, 3.5, 4, 4.5, 5, 5.5, and 6 hours after the dilution; 200 µL of each culture was transferred to a flat-bottom 96-well plate for measurement with a Synergy H1 microplate reader (BioTek Instruments, Inc.). Growth rates were calculated with the following equation:

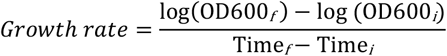

where OD600_*f*_ and Time_*f*_ are from the 3-hour time point data and OD600_*i*_ and Time_*i*_ are from the 1-hour time point data. These time points are selected because results show an exponential growth in this period.

### Animal studies

All mouse experiments were approved by the University of North Texas Institutional Animal Care and Use Committee (Protocol number: 24007) and they were performed in Animal Vivarium of the University of North Texas. All mouse experiments were performed with male 7-to 8-week old C57BL/6 mice (Charles River Laboratories, Wilmington, MA). Mice were housed in a specific pathogen free barrier facility maintained at 30-70% humidity and 68–79 °F under a 12:12 hour light:dark cycle. Mice were provided feed (Purina Conventional Mouse Diet (JL Rat/Mouse 6 F Auto) #5K67) and water ad libitum. A total of 12 mice were used and they were housed in 3 cages, with 4 mice per cage. Among these mice, 4 of them in a cage were served as controls and they lived under these conditions throughout the experiments. The other 8 mice were treated as described below.

After quarantining mice for a week, on Day 1 of the experiment, an amount of 20 mg streptomycin sulfate in 100 µL filter-sterilized water was administered to each mouse by oral gavage using 20G x 30mm plastic feeding tubes, which aims to reduce the native microbiome for enhancing *EcN* colonization. Standard drinking water was switched to filter-sterilized water with 50 g/L sucrose, 5 g/L cellobiose, 100 mg/L ampicillin, and 50 mg/L kanamycin. This modified drinking water is expected to create a permissive environment in the gut by continuously supplying cellobiose. The supplements with ampicillin and kanamycin is expected to further enhance *EcN* colonization as our engineered cells have resistance to these antibiotics but not for most of the native microbes. On Day 3, mice were gavaged with 5×10^8^ CFU *EcN* in 100 μL phosphate buffered saline (PBS). On Day 4, drinking water was switched to filter-sterilized water with 50 g/L sucrose, 5 g/L cellobiose, and 100 mg/L ampicillin. Kanamycin was removed from the water supply such that a loss of the *mcherry*-containing plasmid, which caused a loss of kanamycin-resistance, would not cause stress or death to the *EcN*. On Day 6, drinking water was further switched to create non-permissive and permissive conditions. For mice in the cage that served as the non-permissive group, they were provided with filter sterilized water without cellobiose (only 50 g/L sucrose and 100 mg/L ampicillin); mice in the other cage (permissive group) were continually provided the same drinking water (with 50 g/L sucrose, 5 g/L cellobiose, and 100 mg/L ampicillin).

Drinking water was prepared freshly and replaced daily during the experiment. Fecal samples were collected once per day from Days 4 to 10. At the end of the experiment, mice were sacrificed through carbon dioxide asphyxiation and samples were collected from the cecum (see **Supplementary Figure 6**).

### Preparation of fecal and cecal samples for flow cytometric and CFU analyses

Fecal samples were collected once per day on Days 4 to 10 for *EcN*-treated mice and they were collected once on Day 10 from control mice. Control mice were not treated with *EcN* nor any antibiotics; they lived in standard conditions during this experiment. Cecal samples were collected from all 12 mice after they were sacrificed. Our sample preparation method is modified from a previous study (*31*). Per 5 mg of a fecal or cecal sample, it was resuspended in 1 mL of PBS with 5 g/L cellobiose. The solution was filtered with a 5-µm syring filter to remove particulates and large cells. Filtered samples were incubated at 37 oC for 1 hour, facilitating mCherry protein to obtain oxygen for chromophore activation (*48*). These samples were used immediately for CFU analysis and stored at 4 oC before flow cytometric analysis.

For CFU analysis of mouse samples, filter samples were serially diluted from 10^0^ to 10^7^ folds with PBS. A volume of 5 µL of each diluent was spotted on an LB agar plate with kanamycin, ampicillin, and cellobiose for colonies to form.

### Flow cytometry

A NovoCyte 3000VYB flow cytometer (Agilent Technologies, Inc.; Santa Clara, CA) was used to analyze fluorescence levels in bacterial cells to generate data in **Figures 1B, 1D, 1E, 2A, 2B, 3C, 4B, and 4C** as well as in **Supplementary Figures 1, 2, 3, 5, and 6**. The Novo-Express software (Agilent Technologies, Inc.) was used to perform data analysis. Flow cytometry data were gated by forward and side scatter to eliminate multi-cell aggregates; our data collection settings were to acquire 50,000 counts within these gates or use 20 µL of the sample. Samples were cell cultures (for in vitro experiments) or filtered fecal and cecal samples (for mouse experiments) with a 100-fold dilution (1.5 µL of a culture sample into 150 µL final volume) with phosphate-buffered saline (PBS). The PE-Texas Red channel was used to measure mCherry fluorescent signals and the geometric means of mCherry fluorescence distributions were calculated with the software (**Figure 1D**). To determine the portion of cell population that were expressing mCherry protein, threshold value for mCherry were set based on the background fluorescence intensity of cells without the *mcherry* gene (<10^3^ A.U.; see **Supplementary Figure 1**). The percentage and number of counts with fluorescence values above the threshold value were considered as the portion of cells that were mCherry-active.

### Real-time PCR

A QuantStudio 3 Real-Time PCR system (ThermoFisher Scientific; Waltham, MA) was used to quantify the *mcherry*-containing plasmid in samples. The VWR® qPCR Master Mix kit was used for performing reactions following the manufacturer’s protocol. To generate the standard curve as shown in **Supplementary Figure 4**, a range of quantities of the *mcherry*-containing plasmid (0.002 to 6 ng) was mixed with the biocontainment plasmid to reach a total quantity of 10 ng; these standard samples were used to perform real-time PCR with the two primers as shown in **Supplementary Table 1**, which complement the *mcherry* gene. Four technical replicates were performed for each standard and the average was used for generating the standard curve. A linear equation was obtained for the threshold cycles (C_t_) values versus the log_10_ values of *mcherry*-containing plasmid quantities. To determine quantities of the *mcherry*-containing plasmid in engineered cells, plasmids from cell samples were isolated with a Miniprep kit (Qiagen, Germany). A quantity of 10 ng of each plasmid sample was used for real-time PCR; four technical replicates were performed for each biological sample and the averaged C_t_ value was fit into the equation from the standard curve to determine plasmid quantity. Data were plot and statistically analyzed with GraphPad Prism 7.05.

### Data presentation and statistical analysis

Data were plotted and statistically analyzed with GraphPad Prism 7.05. For comparing results with background noise levels (**Figures 1E, 2B, 3C, 4B, 4C, and Supplementary Figure 7**), the log value of data from each biological replicate was calculated, which was then used for two-tailed unpaired t-test analysis to compare with control data.

For evaluating the standard curve of RT-PCR experiments (**Supplementary Figure 4**), linear regression analysis was performed to gain the R^2^- and P-values.

